# EyeHex toolbox for complete segmentation of ommatidia in fruit fly eyes

**DOI:** 10.1101/2024.03.04.583398

**Authors:** Huy Tran, Nathalie Dostatni, Ariane Ramaekers

## Abstract

Variation in *Drosophila* compound eye size is studied across research fields, from evolutionary biology to biomedical studies, requiring the collection of large datasets to ensure robust statistical analyses. To address this, we present EyeHex, a tool for automatic segmentation of fruit fly compound eyes from brightfield and scanning electron microscopy (SEM) images. EyeHex features two integrated modules: the first utilizes machine learning to generate probability maps of the eye and ommatidia locations, while the second, a hard-coded module, leverages the hexagonal organization of the compound eye to map individual ommatidia. This iterative segmentation process, which adds one ommatidium at a time based on registered neighbors, ensures robustness to local perturbations. EyeHex also includes an analysis tool that calculates key metrics of the eye, such as ommatidia count and diameter distribution across the eye. With minimal user input for training and application, EyeHex achieves exceptional accuracy (>99.6% compared to manual counts on SEM images) and adapts to different fly strains, species, and image types. EyeHex offers a cost-effective, rapid, and flexible pipeline for extracting detailed statistical data on *Drosophila* compound eye variation, making it a valuable resource for high-throughput studies.

## Background

Insect compound eyes consist of crystal-like lattices of elementary eyes, called facets or ommatidia. Each ommatidium is constituted of a fixed number of cells, including photoreceptors that sense light and transmit visual information to the brain, as well as a set of accessory cells (Nériec and Desplan, 2016; Wolff and Ready, 1993). Compound eye size varies greatly among insect groups and species due to differences in the number or diameter of individual facets. In fruit flies (*Drosophila* genus), the number of ommatidia per eye ranges from approximately 600 to 1500, while ommatidia diameter varies from approximately 15 to 22 μm (Arif et al., 2013; Gaspar et al., 2019; Keesey et al., 2019; Posnien et al., 2012; Ramaekers et al., 2019; Reis et al., 2020a). Genetic variation is the primary driver of eye size variation among fruit flies, but environmental conditions such as food quality, larval density and temperature also play a role (Currea et al., 2018). On a wider phylogenetic range, the variation in eye size is much more spectacular. For example, the number of ommatidia varies from zero in some ant species for instance, up to approximately 20.000 in dragonflies (Land and Nilsson, 2012).

Variation in compound eye size is investigated from various scientific perspectives, ranging from evolutionary studies to biomedical research. Evolutionary ecologists aim to identify how eye size variation affects vision and visually guided behaviors, potentially resulting in adaptation to novel ecological niches. Research has shown that larger eyes account for improved vision in dim light or acuity (reviewed in (Land, 2005; Land and Nilsson, 2012; Warrant, 2008; Warrant and Nilsson, 1998). In contrast, reduced eye size is common in insects adapted to darkness, such as cave-dwelling beetles, which rely little or not at all on vision (Luo et al., 2019, 2018).

Variation in compound eye size, especially in fruit flies, is also of interest to developmental or evolutionary-developmental biologists who investigate the genetic and developmental origins of morphological variation (reviewed in (Casares and McGregor, 2020)). Furthermore, it serves as a genetic model for human eye conditions, such as aniridia or retinitis pigmentosa (reviewed in (Gaspar et al., 2019)) and a “test-tube” for studying various human pathologies (e.g. Zika virus pathogenicity (Harsh et al., 2020) or molecular determinants of spinal muscular atrophy (Maccallini et al., 2020)).

Until now, the development of larger scale studies has been hindered by the lack of a simple, accurate, and cost-effective method to fully segment ommatidia. In most reports, ommatidia were manually counted from Scanning Electron Microscopy (SEM) images with limited throughput (Arif et al., 2013; Keesey et al., 2019; Posnien et al., 2012; Ramaekers et al., 2019). Statistical analyses that requires larger datasets were performed using either estimations of ommatidia numbers (Ramaekers et al., 2019) or measurements of eye surface area or length instead of ommatidia counts (Arif et al., 2013; Harsh et al., 2020; Norry and Gomez, 2017). In a recent report, researchers applied a commercial segmentation tool (Amira v.2019.2, Thermo Fischer Scientific) to high-quality images obtained by X-ray tomography (Gaspar et al., 2020). Brightfield imaging could be a desirable alternative to the more laborious and expensive SEM or X-ray tomography technologies. However, due to the prominent curvature of compound eyes, segmenting ommatidia from 2D images is non-trivial. Two features present specific challenges. The first feature is the diverse degree of ocular hair (named bristles) covering ommatidia in 2D eye images. Depending on their location on the eye surface, the level of coverage varies from minimal (e.g., Fig. 1, black frames) to substantial (e.g., Fig. 1, green, blue and orange frames). The second feature is the dome-shaped transparent corneas, which act as lenses overlaying the ommatidia. This causes the ommatidia to reflect light differently depending on their orientation and the angle of the incoming light, hence the varying levels of reflection observed in brightfield images (Fig. 1A). A recent study on eye size variation in the *Drosophila virilis* group exploited this property, capturing and segmenting the reflection of light by each ommatidia, rather than imaging the ommatidia themselves (Reis et al., 2020a).

**Figure 1.**
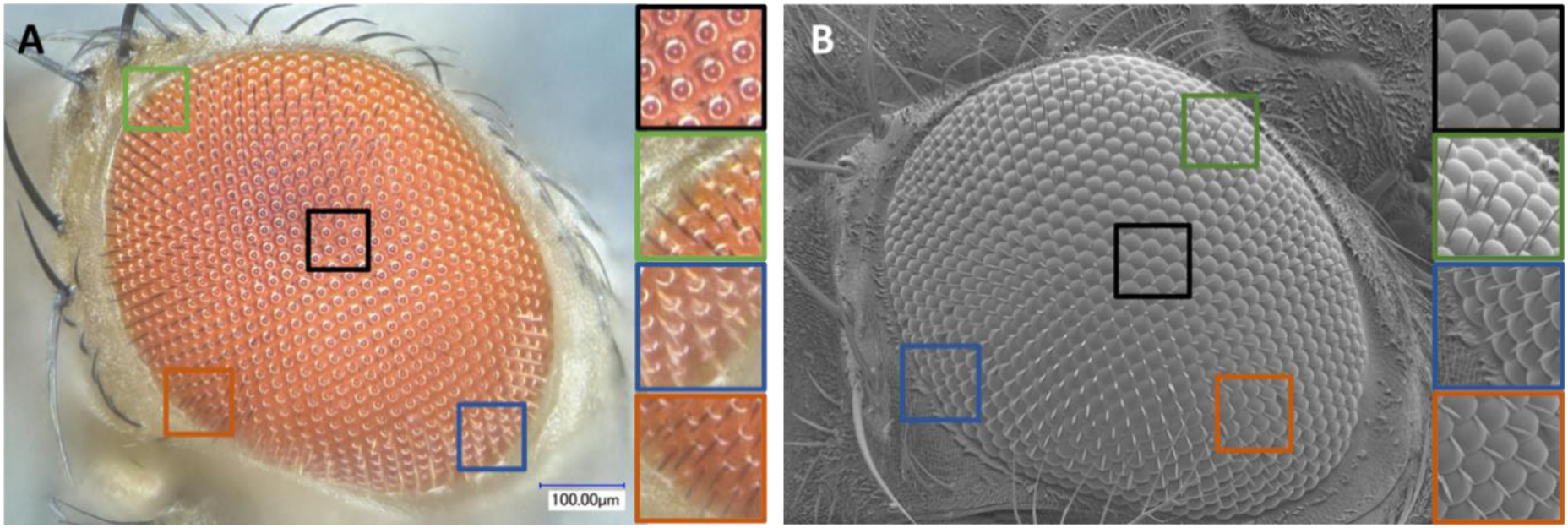
The many facets of the eye. Due to the convex structure of the compound eye, the aspect of ommatidia observed on 2D images varies greatly in different eye regions. Images shown here were acquired using (A) a macroscope (this study) and (B) by SEM (Ramaekers et al., 2019). The zoom-in regions are shown on the right side of each panel with the corresponding frame colors. Note the differences in reflection, shape and inter-ommatidial hair aspect.

Another recent method, named Ommatidia Detection Algorithm (ODA) (Currea et al., 2023), applies Fourier Transform to detect ommatidia on images of glue eye molds, light microscopy and SEM images of various species of insects. This method detects the periodic patterns of ommatidia in the compound eyes with filters in the frequency domain of the images. However, when applied to 2D images, these two hard-coded methods are limited by the high variability of the ommatidia imaging angles in distinct regions of the compound eye (Fig. 1), particularly at the edges where ommatidia are imaged at markedly slanted angles. This may result in the misdetection of ommatidia or the misidentification of ommatidia’s centroid locations.

In this study, we present EyeHex, a comprehensive toolbox for the automated segmentation of ommatidia in *Drosophila* compound eyes. EyeHex consists of two modules (Fig. 2): Module 1 uses a machine learning-based classifier (Trainable Weka Segmentation,

**Figure 2.**
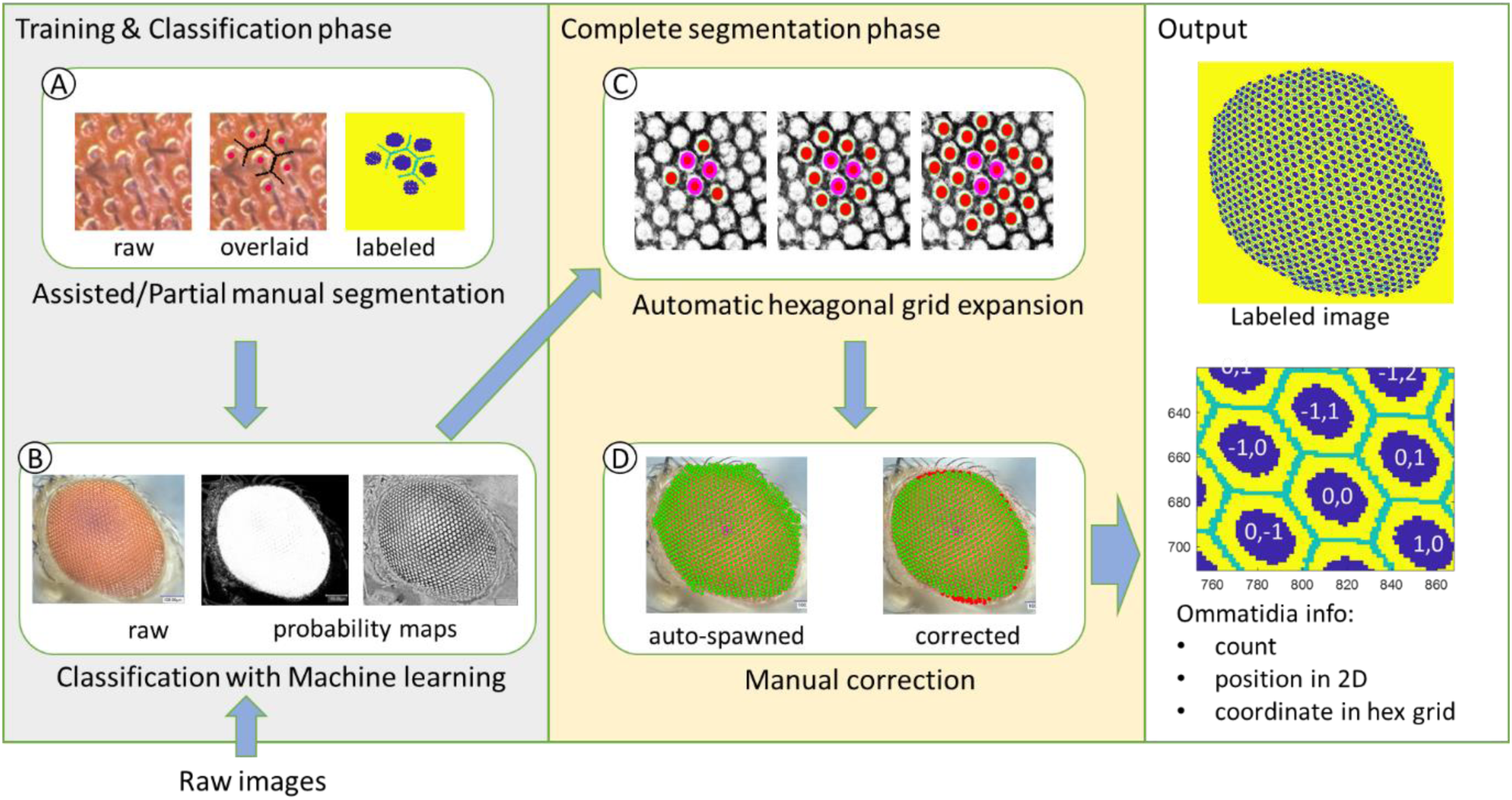
Overview of ommatidia segmentation steps using EyeHex toolbox. The process consists of two phases. The ‘*Training and Classification phase’* involves the generation of the training data with he assisted segmentation of a subset of ommatidia (module A) for the machine learning module, which converts raw input images into probability maps of the eye region and facet region (module B). In the *Complete segmentation phase’*, ommatidia are automatically and exhaustively segmented by expanding a hexagonal grid to fit the facet probability map within the eye region (module C). The user then validates and manually corrects this ommatidia grid using a provided graphical user interface (module D). The outputs include the labelled images for the ommatidia facet and boundary regions (top right), the ommatidia count and the location of each individual ommatidium (bottom right).

ImageJ (Arganda-Carreras et al., 2017) to isolate the compound eye from the background and delineate individual ommatidia. Users can customize training data, enabling broad compatibility with diverse image types, from brightfield and SEM images, acquired using various settings. Module 2 is hard-coded and leverages the hexagonal organization of ommatidia to precisely map their location and quantify their number. This module requires minimal user input, ensuring cost-effective and efficient characterization of *Drosophila* eye size variations. The pipeline is coupled with an analytical tool that automatically generates high-precision metrics, such as ommatidia counts along the eye anterior-posterior axis and ommatidia diameter distribution across the eye.

## Methods

### Data acquisition

#### *Drosophila* strains and culture

Fly stocks were cultured on standard cornmeal diet food at 25°C in density-controlled conditions: batches of 10 females and 10 males were raised at 25°C and transferred into a fresh vial every day. For each vial, eye measures were performed only on females eclosing during the first two days of eclosion. Genotypes used in this study were: for *Drosophila melanogaster*: Canton-S BH (Ramaekers et al., 2019); Hikone-AS (Kyoto DGRC 105668); DGRP-208 (WT25174; BDSC 25174); w;;;+/Df(4)J2 (abbreviated as WTJ2; Ramaekers et al., 2019); for *Drosophila pseudoobscura*: Catalina Island, California, Isofemale line (National Drosophila Species Stock Center, 14011-121.121).

#### Scanning electron microscopy (SEM) images

SEM images are reused from (Ramaekers et al., 2019).

#### Light-microscopy images (Ramaekers et al., 2019 and this study)

Light macroscopy images were either reused from (Ramaekers et al., 2019; WTJ2) or newly acquired (Hikone-AS). In the latter case, they were acquired from non-fixed, freshly cut adult heads glued laterally on glass slides using a solvent-free all material glue (UHU® - cat #64481). 3D image stacks (Z distance = 8 μm) were acquired at 400-600X magnification with a Keyence digital macroscope VHX 2000 using optical zoom lens VH-Z20R/W. Single focused 2D images were generated using Stack Focuser plugin from ImageJ (Schindelin et al., 2012).

### EyeHex workflow

The EyeHex toolbox protocol consists of two successive steps: (1) Training and Classification, and (2) Segmentation (Fig. 2).

### Automatic compound eye and ommatidia classification with machine learning

#### Training data preparation

This step is required when a new image type (e.g., SEM, brightfield microscope…) is introduced. A first set of 2D sample images is used as training data for the classification of the eye area (vs. non-eye area) and the ommatidia surfaces (vs. ommatidia boundaries) (Module A, Fig. 2). Using the Graphical User Interface (GUI) included in the EyeHex toolbox, users are invited to manually delineate the compound eye area and segment small groups of ommatidia at different locations of the eye (e.g., center vs. edges), thus presenting different shapes, reflections and bristle coverage (Fig. 1A). The machine learning module then uses this training data to convert each raw input image into probability maps of the eye area and ommatidia surfaces (module B, Fig. 2). Provided that the imaging settings are unperturbed, a single training image is sufficient to achieve accurate classification.

#### Automatic classification

Classification is performed using the Fiji Weka plug-in (Arganda-Carreras et al., 2017) through a provided macro. Training datasets are used to train two separate pixel-based Fast Random Forest classifiers, each containing 100 decision trees (Breiman, 2001). The trained classifier will then convert the focused 2D images into probability maps of the eye area and the ommatidia surfaces. The compound eye region is defined by thresholding the eye probability map with the Otsu algorithm (Otsu, 1979).

### Automatic segmentation

EyeHex automatic segmentation takes advantage of the remarkably robust hexagonal organization of the compound eye. During this step, ommatidia will be automatically and comprehensively segmented by expanding a hexagonal grid to fit the ommatidia probability map within the eye area (module C, Fig. 2). Users can validate and manually correct the segmentation process for each image using the provided GUI (module D, Fig. 2). The output of the segmentation consists of (1) a labeled image carrying the information on the location of ommatidia surfaces and boundaries, (2) the count of ommatidia number and (3) data on each ommatidium (2D position and coordinates on the hexagonal grid).

#### The ommatidia hexagonal grid

EyeHex segmentation module maps each ommatidium to a unique coordinate on a hexagonal grid. Thus, each ommatidium is characterized by two sets of coordinates: an integer coordinate (*x*_*hex*_, *y*_*hex*_) on the hexagonal grid and a continuous Cartesian coordinate (*x*, *y*) on the 2D ommatidia probability map (Fig. 3A). Note that each ommatidium of the hexagonal grid shares edges with 6 neighbors, instead of 4 as in a normal orthogonal grid. The segmentation process includes three steps:

**Figure 3.**
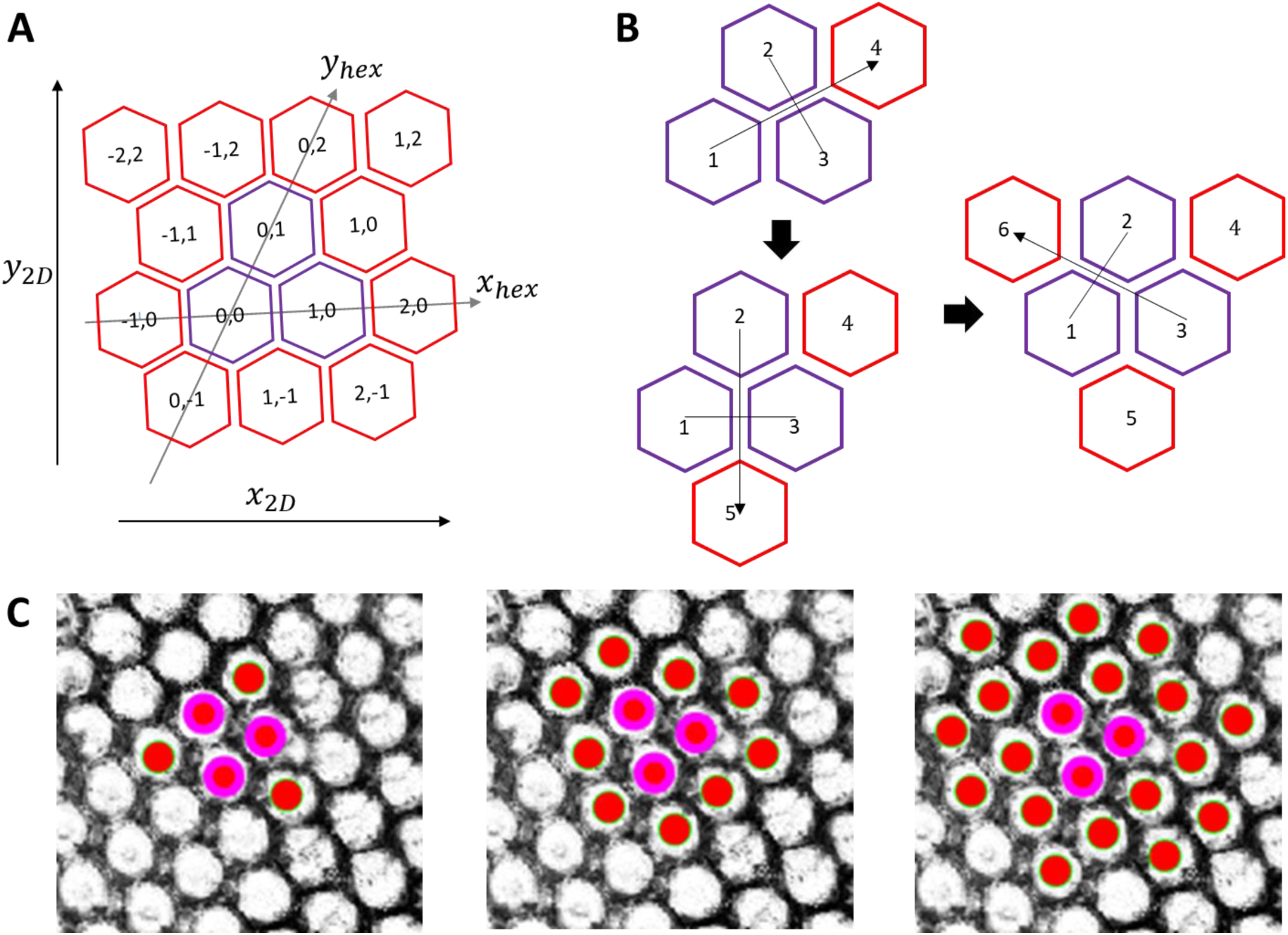
The ommatidia hexagonal grid. (A) The grid is seeded from three original ommatidia (purple). Each ommatidium is identified by a unique integer coordinate (*x*_*hex*_, *y*_*hex*_) on the hexagonal grid, alongside its continuous Cartesian coordinate (*x*_2𝐷_, *y*_2𝐷_) on the ommatidia probability map. Note that the hexagonal axes and the Cartesian axes are not necessarily parallel. (B) New ommatidia (red) are added individually to the grid by mirroring the seed ommatidia (purple): ommatidium 4 is spawned by mirroring ommatidium 1 through ommatidium 2 and 3; ommatidium 5 by mirroring ommatidium 2 through ommatidium 1 and 3 and ommatidium 6 by mirroring ommatidium 3 through ommatidium1 and 2. (C) New ommatidia (red solid circles) are segmented and added to the outer row of existing hexagonal grid set by 3 seed ommatidia (purple), shown here on top of the ommatidia probability map 𝐼_𝑝𝑟𝑜𝑏_ generated during classification phase.

Step 1: Setting the origin and orientation of the hexagonal grid:

The origin and orientation of the grid are established based on three original ommatidia adjacent to one another (purple cells in Fig. 3A; with coordinates (*x*_*hex*_, *y*_*hex*_) equal to (0,0), (1,0), (0,1)), selected by the user at the center of compound eye.

Step 2: Expansion of the hexagonal grid:

The hexagonal grid is expanded by adding individual ommatidia to the current grid. For each three adjacent ommatidia (referred as omtd1, omtd2, omtd3; purple cells in Fig. 3B), we have their respective coordinates on the hexagonal grid (*x*_*hex*1_, *y*_*hex*1_), (*x*_*hex*2_, *y*_*hex*2_), (*x*_*hex*3_, *y*_*hex*3_) and on the Cartesian grid (*x*_1_, *y*_1_), (*x*_2_, *y*_2_), (*x*_3_, *y*_3_). Those coordinates are used to predict the position of a fourth ommatidium, omtd4, which mirrors omtd1 through omtd2 and omtd3 (Fig. 3B) via Eq. 1-4:

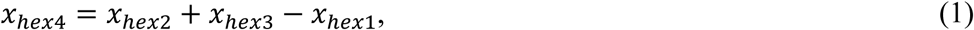

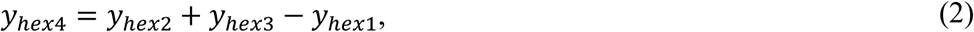

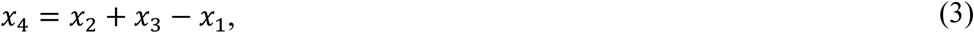

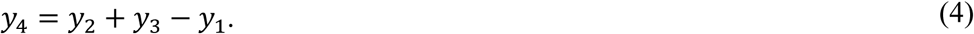

Based on the ommatidia probability map, the program will find an ommatidium within the vicinity of the mirrored position (*x*_4_, *y*_4_) and add omtd4 to the hexagonal grid at the coordinates (*x*_*hex*4_, *y*_*hex*4_) (see below). Only the closest ommatidium within a distance of 𝐿_𝑔𝑟*i*𝑑_/2 from the predicted position (*x*_4_, *y*_4_) is added to the grid. Here, 𝐿_𝑔𝑟*i*𝑑_is the average length of the hexagonal ommatidial edges:

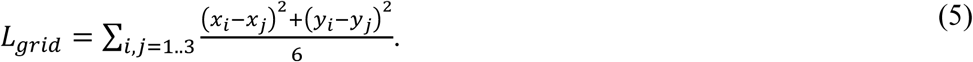

During the automatic hexagonal expansion (Fig. 3C), new ommatidia close to the mirrored position that best fits the ommatidia facet probability map (see below) will always be added to the perimeter of the grid, unless they are found outside the compound eye region (Fig. 3C).

#### Fitting new ommatidia to the ommatidia probability map

From the three adjacent ommatidia (omtd1, omtd2, omtd3), we look for the position of omtd4 (*x*_4_, *y*_4_) based on the facet probability map (denoted as 𝐼_𝑝𝑟𝑜𝑏_(*x*, *y*)) generated with the machine learning module. The general strategy is “find Mickey pattern” with omtd2 and omtd3 being the “ear” and omtd4 being the “face”. A template image 𝐼_𝑚𝑜𝑢𝑠𝑒_ of size 𝑚 × 𝑚 (𝑚 in pixel) is constructed with three filled identical circles of radius 0.225× 𝑚, the centers of which form a perfect angle with edge 0.5 × 𝑚 (Fig. 4A). The pattern is then smoothened with a 2D gaussian filter with 𝜎=𝑚/20. We also set an “ignore” region (with *NaN* pixel value, dashed blue in Fig. 4A) on 𝐼_𝑚𝑜𝑢𝑠𝑒_ to avoid interference from surrounding ommatidia.

**Figure 4.**
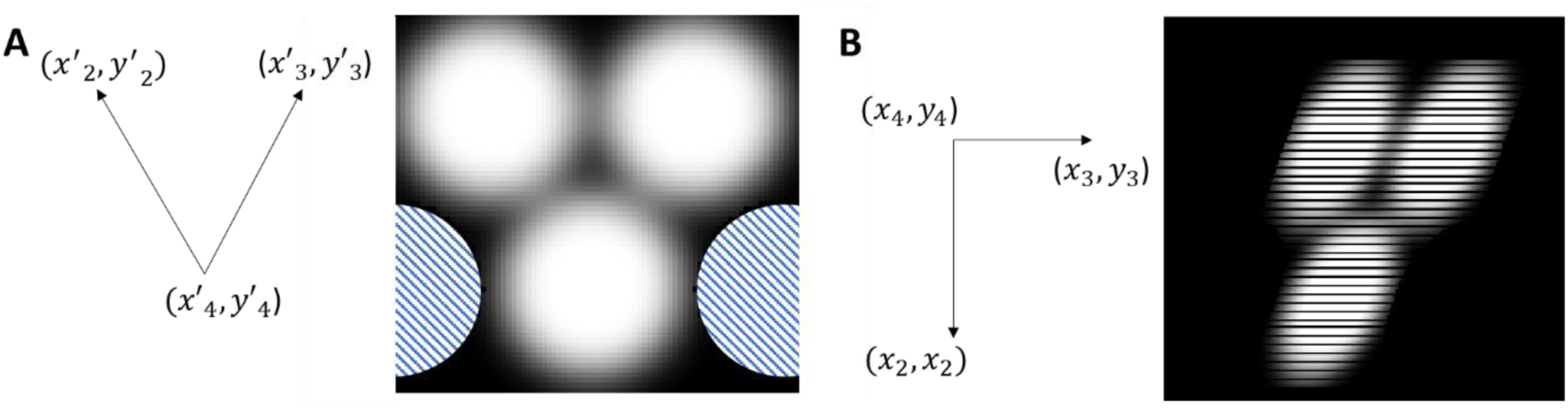
‘*Mickey patterns*’ on the facet probability map. (A) The coordinate vectors 𝐵′ =(𝑣′_𝑎_,𝑣′_𝑏_) (left) for the mouse image 𝐼_𝑚𝑜𝑢𝑠𝑒_ (right). The objective function does not consider the blue dashed regions (NaN value). (B) Example of the new coordinate vectors 𝐵 =(𝑣_𝑎_,𝑣_𝑏_) from ommatidium 2, ommatidium 3 and ommatidium 4 position (left) with the transformed 𝐼_𝑚𝑜𝑢𝑠𝑒_ in this coordinate (right).

We called (*x*_2_′, *y*^′^_2_), (*x*^′^_3_, *y*^′^_3_) and (*x*^′^_4_, *y*^′^_4_) the coordinates of the centers of the three circles in the template image 𝐼_𝑚𝑜𝑢𝑠𝑒_. We find a linear transformation 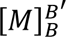 from the coordinate vectors 𝐵′ to the coordinate vectors 𝐵, in which:

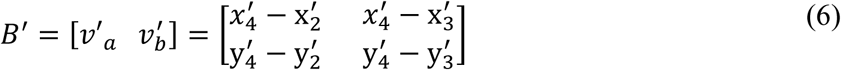

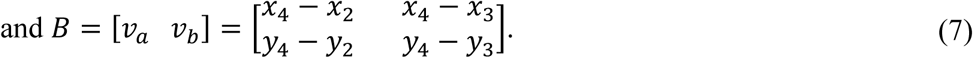

For each non-*NaN* pixel in position (*i*, *j*) in 𝐼_*mouse*_, we find the corresponding coordinate 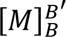 (*i*, *j*) on 𝐼_*prob*_ (Fig. 4B).

The objective function for the fit of the new omtd4 position Δ(*x*_4_, *y*_4_) is given by the squared error between the facet probability map and the transformed mouse image:

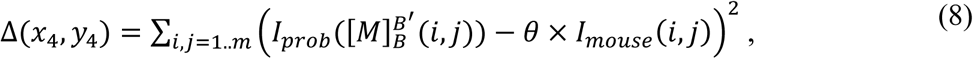

with 𝜃 being a free parameter to adjust for pixel brightness.

Due to frequent irregularities on the probability map, we introduce additional constraints to preserve the integrity of the hexagonal grid and account for the fact that the facet orientation changes gradually along the grid: The objective function Δ(*x*_4_, *y*_4_) is set to infinity when the new omtd4 position is farther than 0.3 × 𝐿_𝑔𝑟*i*𝑑_ from the predicted position that mirrors omtd1, and when the omtd4 top view angle is too steep (<20°) or too different from that of omtd1.

### Manual correction

Following the automatic hexagonal grid expansion, the user can manually remove or add ommatidia using the provided GUI. After manual correction, newly added ommatidia are mapped and automatically given their own coordinate in the existing hexagonal grid.

### Data analyses

#### Measurement of ommatidia size

When multi-focused brightfield images are used, the z-coordinate of each ommatidium can be determined from its position in the original image stack. We use this information to estimate the size of an ommatidium by averaging its Euclidean distances in the three-dimensional space to its adjacent ommatidia. These values are fitted to a polynomial surface to generate a smooth profile of ommatidia altitude and spacing across the eye, as shown in Figure 5B and 5C.

**Figure 5.**
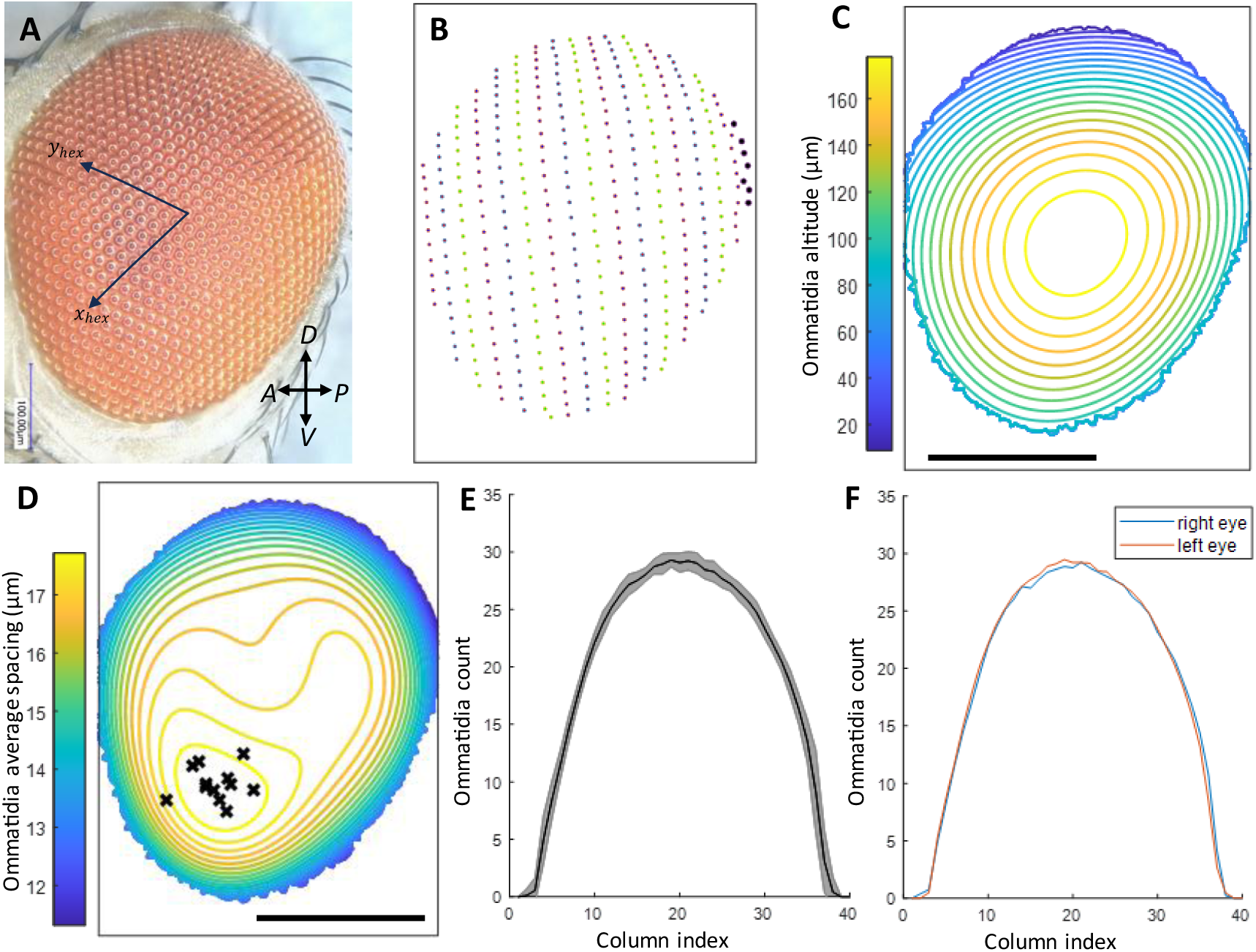
EyeHex output: Quantitative analysis of compound eye features. **A)** Focused brightfield image of a Hikone-AS eye, with the overlayed hexagonal grid axes. **B)** Detected ommatidia centers, sorted into columns (separated by color). Ommatidia in the most posterior (rightmost) column are highlighted in bold. **C)** Contour map of ommatidia altitude (in μm, calculated relative to the lowest focal plane). **D)** Contour map of the average spacing between adjacent ommatidia (in μm) projected onto the eye in panel A. Black crosses indicate the locations of greatest inter-ommatidial distance (largest ommatidia) for individual Hikone-AS eyes; data from *n*=12 eyes. **E)** Number of ommatidia in each column from posterior (P) to anterior (A) (line and shading represent the mean and standard deviation, respectively, n=12). **F)** Comparison of mean ommatidia count profiles between left (blue curve, n =8) eyes and right (red curve, n=4) eyes. Scale bars in C and D: 200 μm. Eye orientation is shown in panel A.

#### Ommatidia column identification and column profiles

The ommatidia are arranged in columns that reflect the progressive specification of R8 ommatidia founder cells along the anterior-posterior axis of the eye primordium (eye-antennal disc) (Ready et al., 1976). To locate these columns, the origin of the ommatidia hexagonal grid (*x*_*hex*_=0, *y*_*hex*_=0) is shifted to the detected center-most ommatidia. The grid is then reoriented so that both the x- and y-coordinates of the most anterior ommatidium in this grid take positive values (Fig. 5A). The sum of the new xy-coordinates of the ommatidia (*x*_*hex*_+*y*_*hex*_) defines the column index that corresponds to their birth order during R8 specification. This column index increases from the posterior-dorsal to anterior-ventral regions of the eye (example in Fig. 5D). From the reoriented grid, we calculate the “column profile” as the number of ommatidia per column, which provides information about the eye geometry (example in Fig. 5E).

#### Averaging of morphological profiles

We accumulate measurements of individual ommatidia from different eyes based on their unique coordinate in the hexagonal grid (*x*_*hex*_, *y*_*hex*_). This allows us to calculate the average ommatidia spacing in different regions of the grid. For visualization, we project this average spacing profile onto a reference eye, where the Cartesian coordinate for each segmented ommatidia is known (example in Fig. 5D).

### Outputs

After manual correction of segmentation and analysis of each eye, EyeHex automatically exports the label image of the identified ommatidia and eye region. These label images can be added to the training data to improve subsequent classifications. Also exported is a comma-separated-values (.csv) file containing information about each ommatidium’s index, its Cartesian and hexagonal coordinates, and its birth order during R8 specification.

## Results

### The EyeHex toolbox

The EyeHex toolbox is a set of MATLAB scripts that allows users to segment all visible ommatidia from 2D images of *Drosophila* compound eyes in a semi-automatic manner. Limited user attention through Graphical User Interfaces (GUIs) is required during the training and manual correction step. EyeHex output includes labeled eye images depicting the ommatidia masks (Fig. 2E), as well as ommatidia counts and positions in .csv format (Fig. 2F) for further statistical analyses. The toolbox, its manual and sample images are available for download on GitHub at https://github.com/huytran216/EyeHex-toolbox.

### Segmentation on SEM images

The gold standard for quantifying ommatidia numbers relies on manual counts performed on 2D compound eye images acquired via scanning electron microscopy (SEM). To evaluate the efficiency of EyeHex segmentation, we compared the number of ommatidia generated by EyeHex toolbox with manual counts from previously published SEM datasets (Ramaekers et al., 2019) for three *D. melanogaster* strains : Hikone-AS (Fig. S1), Canton-S BH (Fig. S2), and DGRP-208 (Fig. S3). For training datasets, we manually segmented approximately 80 ommatidia (∼ 10% of the total per eye) in a single SEM image. Subsequent steps included classification, automatic segmentation, manual corrections, and counting (Table 1; Suppl. File 2). Compared to manual counts, EyeHex ommatidia numbers deviated by 0 to a maximum of 8 ommatidia per eye, achieving an average accuracy exceeding 99.6 % (Table 1, Suppl. File 2). We further tested EyeHex to SEM images of *D. pseudoobscura*, a different *Drosophila* species with significantly larger compound eyes (*D. pseudoobscura*: ∼1200 ommatidia/eye *vs D. melanogaster*: *∼* 800 ommatidia/eye), and compared the EyeHex results to published manual counts (Ramaekers et al., 2019). For this species, EyeHex also yielded highly accurate counts, with differences ranging from 0 to 3 ommatidia per eye (99.9 % accuracy; Table 1, Fig. S4, Suppl. File 2). These results demonstrate our tool’s applicability beyond *D. melanogaster* to other *Drosophila* species.

**Table 1:** Ommatidia counts on SEM and brightfield images.

| Strain | Image type | Image collection | n | EyeHex | EyeHex | Manual | Ellipse |
| --- | --- | --- | --- | --- | --- | --- | --- |
|  |  |  |  | *Pre-correction* | *Post-correction* |  |  |
| Hikone-AS | SEM | *Ramaekers et al. 2019* | 6 | 722.7 ± 26.2 | 741.2 ± 21.8 | 739.7 ± 21.8* | 764.5 ± 26.7 |
| Canton-S<sup>BH</sup> | SEM | *Ramaekers et al. 2019* | 9 | 948.3 ± 31.4 | 857.8 ± 30.1 | 859.3 ± 27.4* | 888.2 ± 33.1 |
| DGRP-208 | SEM | *Ramaekers et al. 2019* | 8 | 859.0 ± 82.1 | 754.5 ± 44.1 | 754.6 ± 43.4* | 797.8 ± 48.1 |
| *D. pseudoobscura*<br>Catalina Island | SEM | *Ramaekers et al. 2019* | 5 | 1192.8 ± 14.6 | 1176.2 ± 22.2 | 1175.0 ± 22.1* | 1216.9 ± 22.0 |
| Hikone-AS | Brightfield | This study | 12 | 787.6 ± 18.2 | 752.7 ± 15.6 | / | 778.7 ± 18.3 |
| WTJ2 (white eyes) | Brightfield | *Ramaekers et al. 2019* | 8 | 870.3 ± 73.3 | 789.9 ± 27.2 | / | 819.6 ± 28.7* |

*Ommatidia counts (mean ± standard deviation) for SEM or brightfield images. The images were either reused from a previous study (Ramaekers et al., 2019) or newly collected. We compare ommatidia counts performed using EyeHex segmentation before and after manual correction, together with fully manual counts ('Manual') or elliptic estimations ('Ellipse'). All strains are from D. melanogaster species, except where otherwise indicated. n: sample size; \*: counts from Ramaekers et al., 2019.*

### Segmentation on brightfield images

The sample preparation and image acquisition procedure for SEM images are relatively time-consuming and expensive. To reduce time and costs, we applied EyeHex on brightfield images acquired using a macroscope on non-fixed samples. Our previous work estimated ommatidia numbers in such images by assuming a perfect elliptical fit for the ommatidia hexagonal grid (see Ramaekers et al., 2019 for the method), but direct manual counts were not performed. To assess whether EyeHex achieves accuracy comparable to SEM with brightfield images, we generated training datasets by manually segmenting 150 ommatidia from two newly acquired Hikone-AS brightfield images. We then applied EyeHex for classification, segmentation and counting (Fig. 5A-B and Table 1, Suppl. File 2). Notably, the Hikone-AS strain used here may differ from that in our previous study (Ramaekers et al., 2019), possibly because of genetic drift or slightly different culture conditions, as experiments were conducted in different laboratories over 5 years apart. Despite this, EyeHex-derived counts closely matched previous manual SEM counts (Table 1), indicating that brightfield imaging combined with EyeHex provides a reliable representation of biological samples. In contrast, our earlier elliptical estimation method systematically overestimated the number of ommatidia by 20 to 40 ommatidia on average (Ramaekers et al. 2019 and Table 1). We also tested EyeHex on *white* mutant eyes (strain WTJ2), as this is one of the most commonly used genetic markers in fruit flies. Those eyes lack the red pigmentation and exhibit reduced contrast. While ommatidia segmentation accuracy remained comparable to other datasets, more manual correction was required - up to 14% of the total ommatidia (Table 1, Suppl. File 2). This primarily involved falsely detected ommatidia at the eye boundary, caused by overestimation of the eye region during the classification step. However, ommatidia segmentation performed equally well compared to the other datasets, confirming EyeHex’s adaptability to low-contrast samples.

### Ommatidia size profile and ommatidia column count

Insect compound eyes often exhibit local specializations – named acute or bright zones – that enhance acuity or signal-to-noise ratio in specific regions of the visual field (Horridge, 1978) (Land, 1997). The presence and location of these acute/bright zones correlate with distinct visually guided behaviors. For example, upward-pointing zones are frequently associated to predatory behavior, whereas male-specific forward-pointing zones are linked to sexual pursuit. In many insect species, including flies, these zones are characterized by enlarged ommatidia. In addition to quantifying ommatidia number, EyeHex can automatically detect these regions of enlarged ommatidia by analyzing ommatidia size variation across the eye. By analyzing brightfield multifocus image stacks, EyeHex can reconstruct the 3D position of each ommatidium (Fig. 5C). This, in turn, enables the calculation of the average distance in 3D space between each ommatidium and its 6 adjacent neighbors (see Methods), providing a proxy for ommatidial diameter across the eye. In all eyes analyzed (e.g., Fig. 5D), ommatidial density and diameter varied by up to 20 % along the posterior-dorsal to anterior-ventral axis of the eye, consistent with observations by (Gaspar et al., 2020) and (Buffry et al., 2024) (Fig. 5D and Fig. S7). Projecting density maps from multiple specimens onto a common reference revealed a consistent zone of enlarged ommatidia in the anterior-ventral part of the eye (black crosses in Fig. 5D and Fig. S7). In *Drosophila*, acute zones have been documented in males, where they occupy an antero-dorsal position. These zones are reportedly absent in females, though the existence and location of analogous structures in female eyes remain unclear. Since our study focused on female eyes, further investigation is required to determine whether the observed antero-ventral ommatidial enlargement represents an acute zone in this sex. Notably, at the extreme edge of the eye, ommatidia diameters measured by EyeHex were unexpectedly small (<12 µm). In these regions, some ommatidia adopt an elongated, non-hexagonal shape, and the diameters reported by EyeHex reflect their reduced width. However, identifying the xy- and z-coordinates of highly slanted ommatidia can be challenging, making the determination of inter-ommatidial spacing at the very edge of the eye prone to errors. Thus, such small diameter may not always fully represent true morphological dimensions.

In the compound eye, ommatidia are arranged in regularly spaced columns along the anterior-posterior (A-P) axis of the eye (Fig. 5B). This organization reflects the progressive determination and spacing of ommatidia founder cells (termed R8 cells) along the A-P axis of the eye primordium during the progression of a differentiation wave named ‘*morphogenetic furrow*’ (Ready et al., 1976) (Wolff and Ready, 1993) (for a recent review, see (Warren and Kumar, 2023)). Thus, quantifying the number of A-P columns in adult eyes provides a read-out of the R8 determination process and of its robustness. Here, we used EyeHex to extract this information from a set of brightfield eye images from 12 individuals of the Hikone-AS wild-type genetic background. Each eye consisted of 33.89 ± 1.05 columns (mean and standard deviation). We also extracted ‘*column profiles*’ by plotting the number of ommatidia per column along the A-P axis of the eye (Fig. 5B), thereby providing a simple representation of the eye geometry. We then aligned the 12 column profiles for comparison. The mean aligned column profile from 12 brightfield eye images is shown in Figure 5E. Interestingly, the standard deviation (shaded area of the plot, Fig. 5E) of the mean is on average ∼1.08 ommatidia, with a maximum of 3.73 ommatidia per column. Therefore, the highly inbred Hikone-AS laboratory individuals show very little variation in eye organization and size, indicating that despite its complexity, compound eye development is remarkably robust. Comparison of the mean aligned column profiles between left and right eyes from separate individuals also did not reveal any significant bias (an average difference of 0.45 ommatidia per column) (Fig. 5F).

## Discussion

The EyeHex toolbox automates segmentation and mapping in 2D space of ommatidia in *Drosophila* compound eyes. Coupled with a dedicated analysis pipeline, it quantifies metrics of the eye organization, including total ommatidia number, ommatidia count profiles along the A-P axis, and contour maps of ommatidial diameters. Using a machine learning-based approach, EyeHex adapts to diverse image types, with performance improving as datasets expand. Existing tools for automated ommatidial segmentation vary in their methodologies and limitations. Reis *et al*. (Reis et al., 2020b) used brightfield images to segment ommatidia reflections as discrete cells separated by a predefined minimum distance – a strategy prone to false negatives in case of damaged or non-reflective ommatidia. In Gaspar *et al*. (Gaspar et al., 2020), ommatidia segmentation was hard-coded and restricted to 3D X-ray tomography images. The segmentation method developed by Currea *et al*. (Currea et al., 2023) is applicable to various image types and exploits periodic ommatidia patterns via filtering in the frequency domain (Fourier transforms), fitting hexagonal/orthogonal grids to overlapping regions. While efficient, this semi-global approach may struggle with extreme warping, common at the compound eye boundary. By contrast, EyeHex employs an adaptive, local strategy: ommatidia are identified sequentially and integrated into the existing eye grid based on their three nearest neighbors. This accommodates local distortions in the compound eye surface, including warping at the eye edges, while preserving global hexagonal organization. In addition, because EyeHex can accommodate non-uniform ommatidia spacing, the segmentation process should not be affected by minor disruptions or irregularities in the eye lattice found in some mutants. Similarly, EyeHex is also effective for quantifying eye morphology in non-*melanogaster Drosophila* species, which share conserved eye organization, as shown in this study for large-eyed *D. pseudoobscura*. It is also adaptable to images exhibiting lower contrast, such as *white* mutant eyes. However, mutant eyes with severe structural abnormalities may not be accurately segmented by EyeHex. Similarly, highly bulged eyes (obscuring edges in brightfield image stacks) may require alternative imaging techniques such as μCT scans (Gaspar et al., 2020) or indirect imaging of eye molds flattened by incision (Ramirez-Esquivel et al., 2017).

Manual counts of ommatidia from SEM images ommatidia remain the gold standard for ommatidia enumeration. EyeHex achieves > 99.6 % concordance with this gold standard after minimal manual corrections. Applied to macroscope brightfield images of unfixed samples, it replicates SEM-derived statistics, maintaining high accuracy while streamlining sample preparation and processing. Nevertheless, fully automated segmentation methods cannot achieve 100% accuracy – a critical consideration given interindividual variation in ommatidial counts (relative standard deviation < 2% in inbred lines; this study, (Currea et al., 2018; Gaspar et al., 2020). Thus, user-guided inspection and manual correction, integrated into EyeHex toolbox, remains essential.

Compound eyes, though widespread across the animal kingdom, exhibit lower acuity than other eye types, such as the vertebrate camera eyes (Kirschfeld, 1976; Land, 1997; Mallock, 1894). To achieve human-level resolution, compound eyes should consist of millions of ommatidia, reaching a radius of 6 meters. Many species compensate via specialized zones (“acute” or “bright” zones), often characterized by enlarged ommatidia, which enhance acuity for specific parts of the visual field (Land, 1997). Despite the adaptive relevance of this trait, most *Drosophila* studies report only average or local measures of ommatidial diameter (Gaspar et al., 2020; Keesey et al., 2019; Posnien et al., 2012; Ramaekers et al., 2019). Exceptions include a recent study applying a 3D version of the ODA segmentation tool (Currea et al., 2023) to synchrotron microtomography images (Buffry et al., 2024). Similarly, EyeHex derives 3D altitude data from brightfield multifocus stacks to map ommatidial diameter across the eye surface, enabling identification of potential acute zones or subtle patterns of variation.

Variation in *Drosophila* compound eye size is a broadly used model in evolutionary developmental biology (Casares and McGregor, 2020). Despite well-documented eye size variation, technical limitations have historically limited large-scale studies, precluding the compound eye’s use in addressing fundamental evo-devo questions. For instance, quantifying subtle morphological variation – a critical metric for assessing developmental precision, stochasticity, or robustness – demands expansive datasets. This bottleneck has relegated studies of phenomena like fluctuating asymmetry in *Drosophila* to simpler traits (e.g., wing morphology and mechanosensory hair patterns, (Debat et al., 2009; Polak and Starmer, 2001), which are more readily quantified. The compound eye, with its extensively characterized development, offers an ideal system to explore these questions in a complex tissue, uniquely positioned to link developmental mechanisms with phenotypic outcome. EyeHex bridges this methodological gap by enabling providing accurate segmentation and extraction of eye morphology statistics across imaging platforms, including accessible brightfield macroscopy. Our pipeline - from imaging unfixed heads to EyeHex analysis – offers a rapid, cost-effective solution for large-scale quantification of eye morphology.

## List of abbreviations

A-P: Anterior-Posterior
μCT: Micro Computed Tomography
SEM: Scanning Electron Microscopy
GUI: Graphical User Interface

## Declarations

### Ethics approval and consent to participate

Not applicable

### Consent for publication

Not applicable

### Availability of data and materials

EyeHex toolbox: code and user manual available to download from https://github.com/huytran216/EyeHex-toolbox.

Examples of segmentation results and analysis for SEM images (Hikone-AS, Canton-S, DGRP-208, *Drosophila pseudoobscura*) and brightfield images (Hikone-AS, WTJ2): Supplementary_file1.pdf

Ommatidia counts: Supplementary_file2.xlsx

### Competing interests

The authors declare that they have no competing interests.

### Author’s contributions

AR collected the data (sample preparation and imaging), HT created EyeHex toolbox and analyzed the data, AR and HT interpreted the data. AR, HT and ND drafted the manuscript.

## Acknowledgments

The authors want to thank Virginie Courtier (CNRS, Institut Jacques Monod, Paris) for providing access to the Keyence macroscope.

## Fundings

HT is supported by Research Council of Finland (354987), Tampere University’s Health Data Science Profile. This work was also supported by “Emergence” grant from Sorbonne Université to AR.

## Notes

### Competing Interest Statement

The authors have declared no competing interest.

### Summary of Updates

New data on WTJ2 Drosophila and D. pseudoobscura fly added (Table 1) Fig 5 updated with more detailed morphological analyses Revised results and conclusions on conclusions with existing tools. Supplementary File 2 added

