## Supplementary file 1 for "EyeHex toolbox for complete segmentation of ommatidia in fruit fly eyes"

### 1. EyeHex ommatidia segmentation on Scanning Electron Microscope (SEM) images.

For each example SEM image (**Fig. S1-S4**), we show: (**A**) the original image; (**B**) the probability map of ommatidia location generated by the machine-learning module; (**C**) the anterior to posterior columns of detected ommatidia, separated by color and progressive dimming; (**D**) ommatidia count in each column from posterior to anterior. The origin of the data is indicated in each figure title.

**Figure S1: Hikone-AS** *(*data from *Ramaekers et al., 2019)*

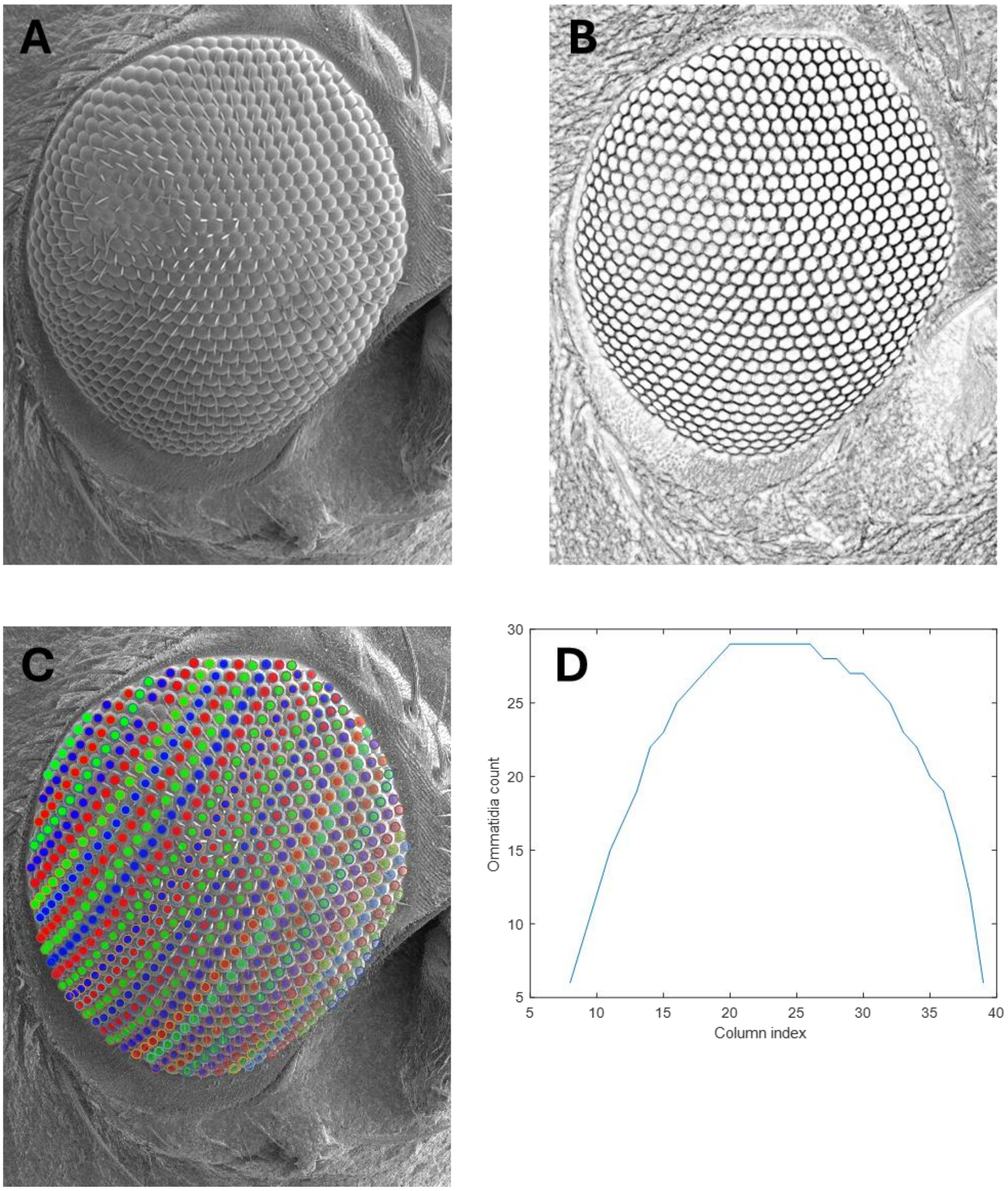


Figure S1. Analysis from EyeHex on a SEM image of a Hikone-AS eye

**Figure S2: Canton-S** *(*data from *Ramaekers et al., 2019)*

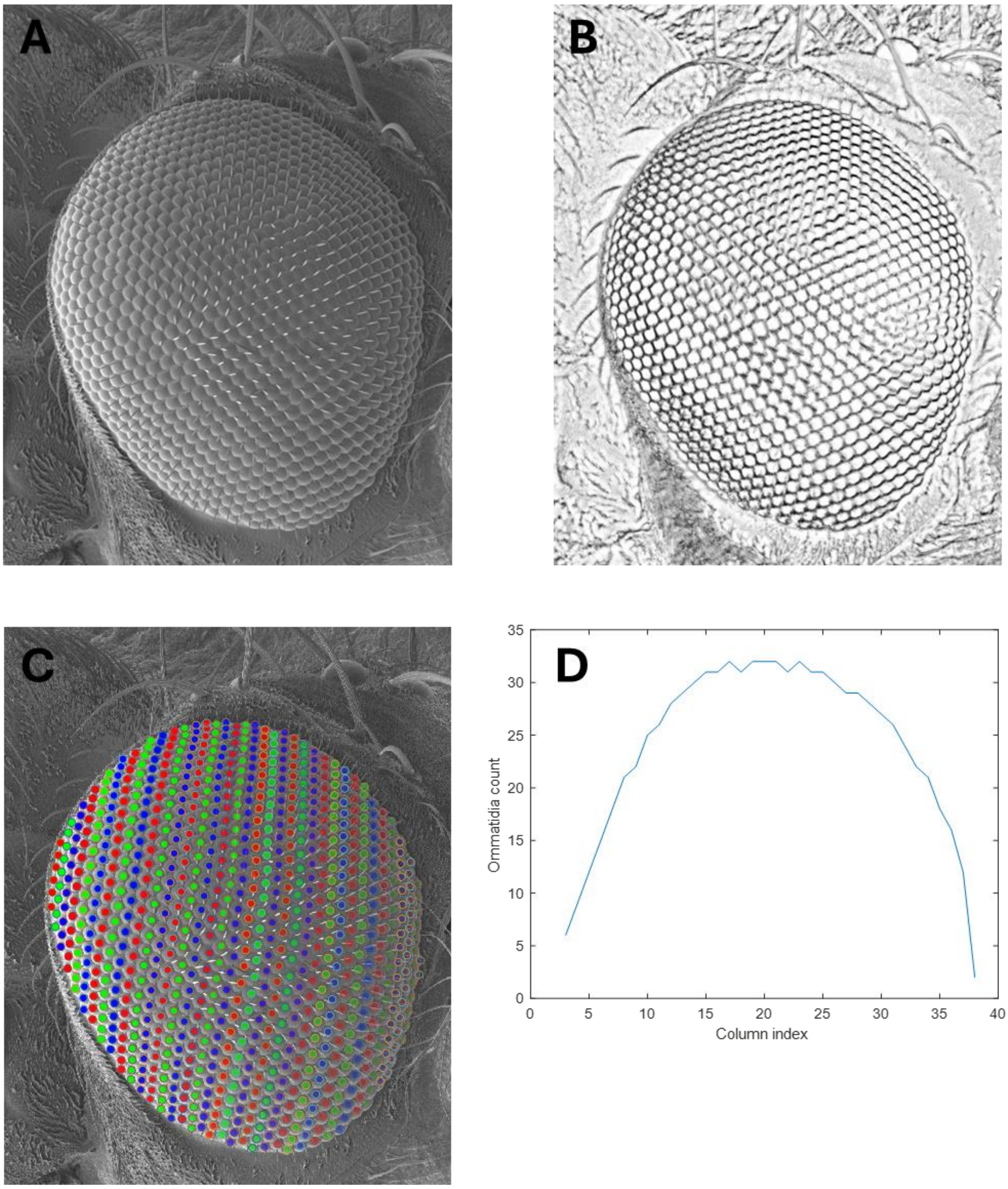


Figure S2. Analysis from EyeHex on a SEM image of a Canton-S eye

**Figure S3: DGRP-208** *(*data from *Ramaekers et al., 2019)*

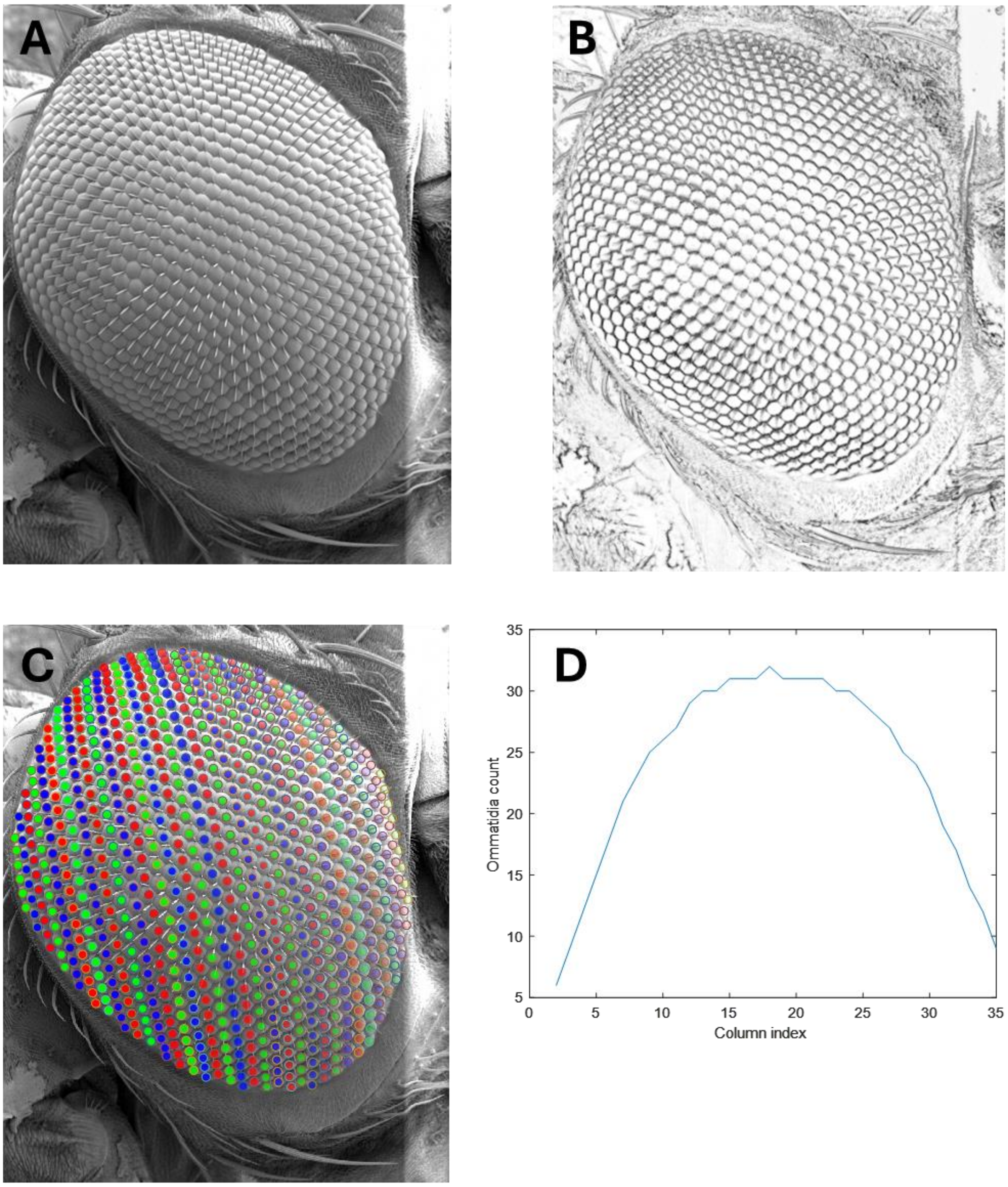


Figure S3. Analysis from EyeHex on a SEM image of a DGRP-208 eye

**Figure S4 : Drosophila pseudoobscura** (data from *Ramaekers et al, 2019*)

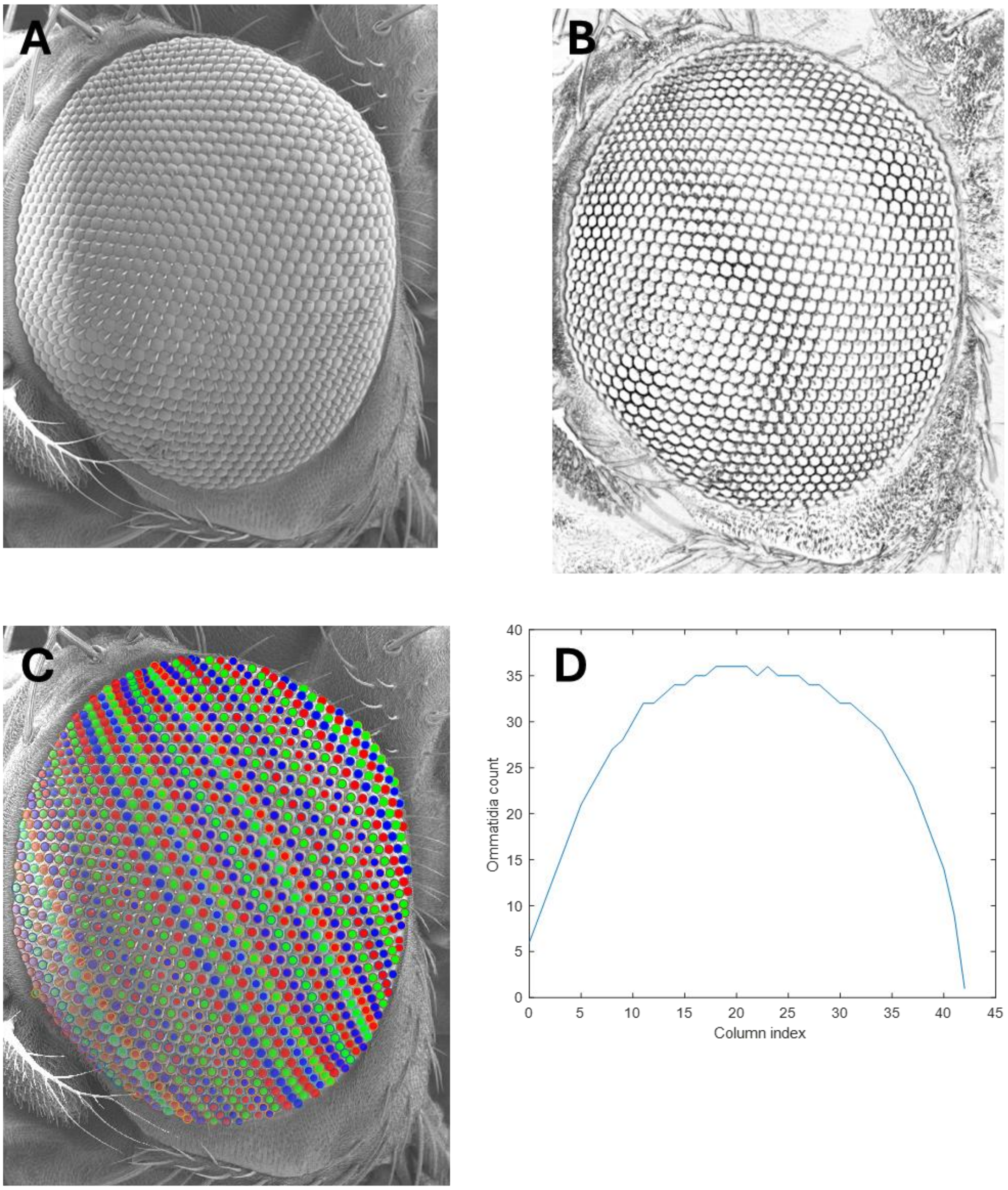


Figure S4. Analysis from EyeHex on a SEM image of a *Drosophila pseudoobscura* eye

#### 2. EyeHex segmentation and morphological analysis on brightfield images.

For each example multi-focus image (Fig. S5-6), we show: (**A**) the focused 2D compound image, scale bar: 200 microns; (B) the probability map of ommatidia location generated by the machine-learning module; (C) the anterior to posterior columns of detected ommatidia, separated by color and progressive dimming; (**D**) a contour map of ommatidia altitude (in micron, calculated relative to the lowest focal plane); (E) a contour map of estimated spacing between individual ommatidia (in micron), (F) ommatidia count in each column from posterior to anterior. The origin of the data is indicated in each figure title.

**Figure S5 : Hikone-AS** (data from this study).

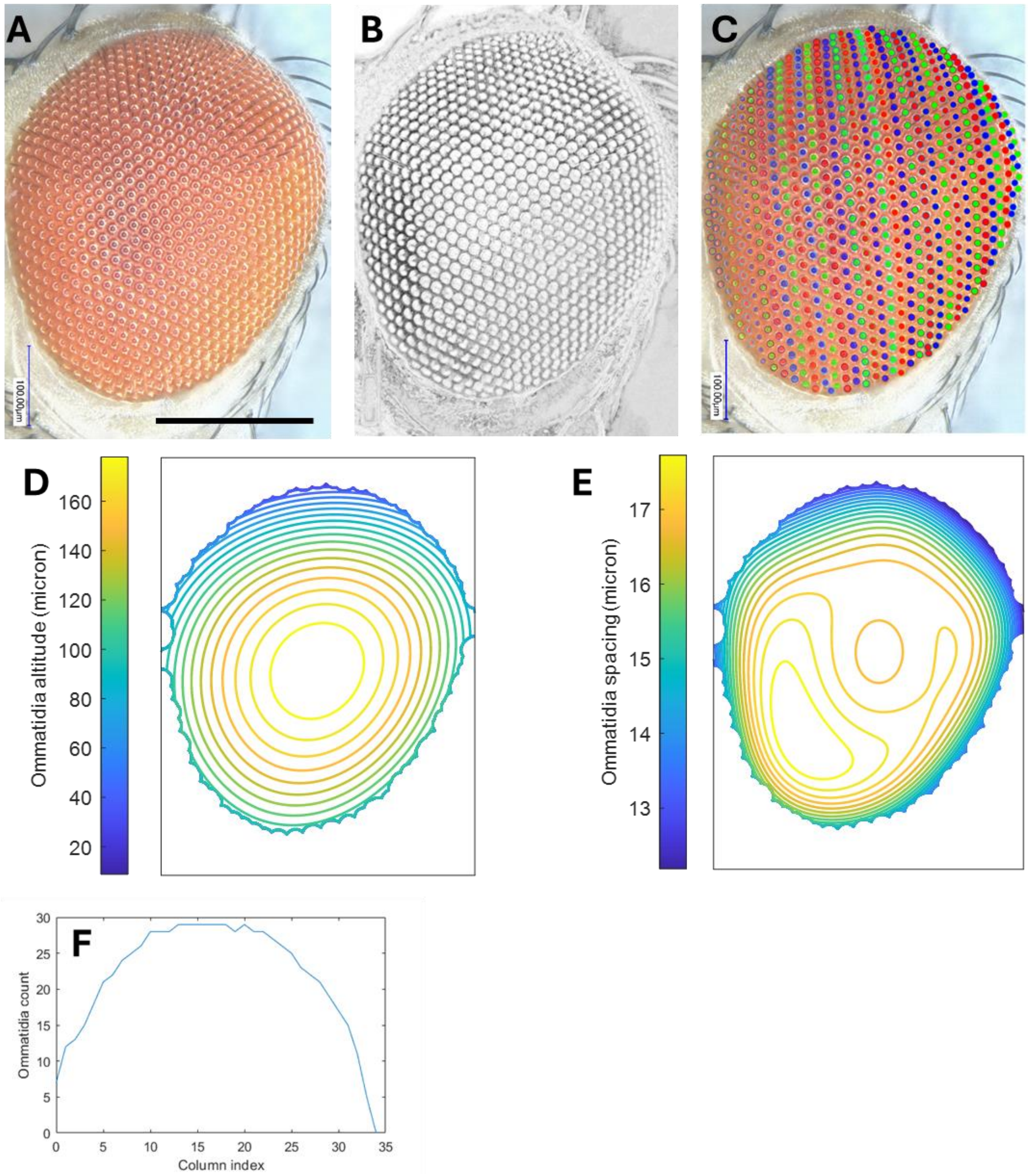


Figure S5. EyeHex morphological analysis on a brightfield image of an Hikone-AS eye

**Figure S6: WTJ2 (F1 progeny from ey3.5G x Df(4)J2)** (data from *Ramaekers et al., 2019*)

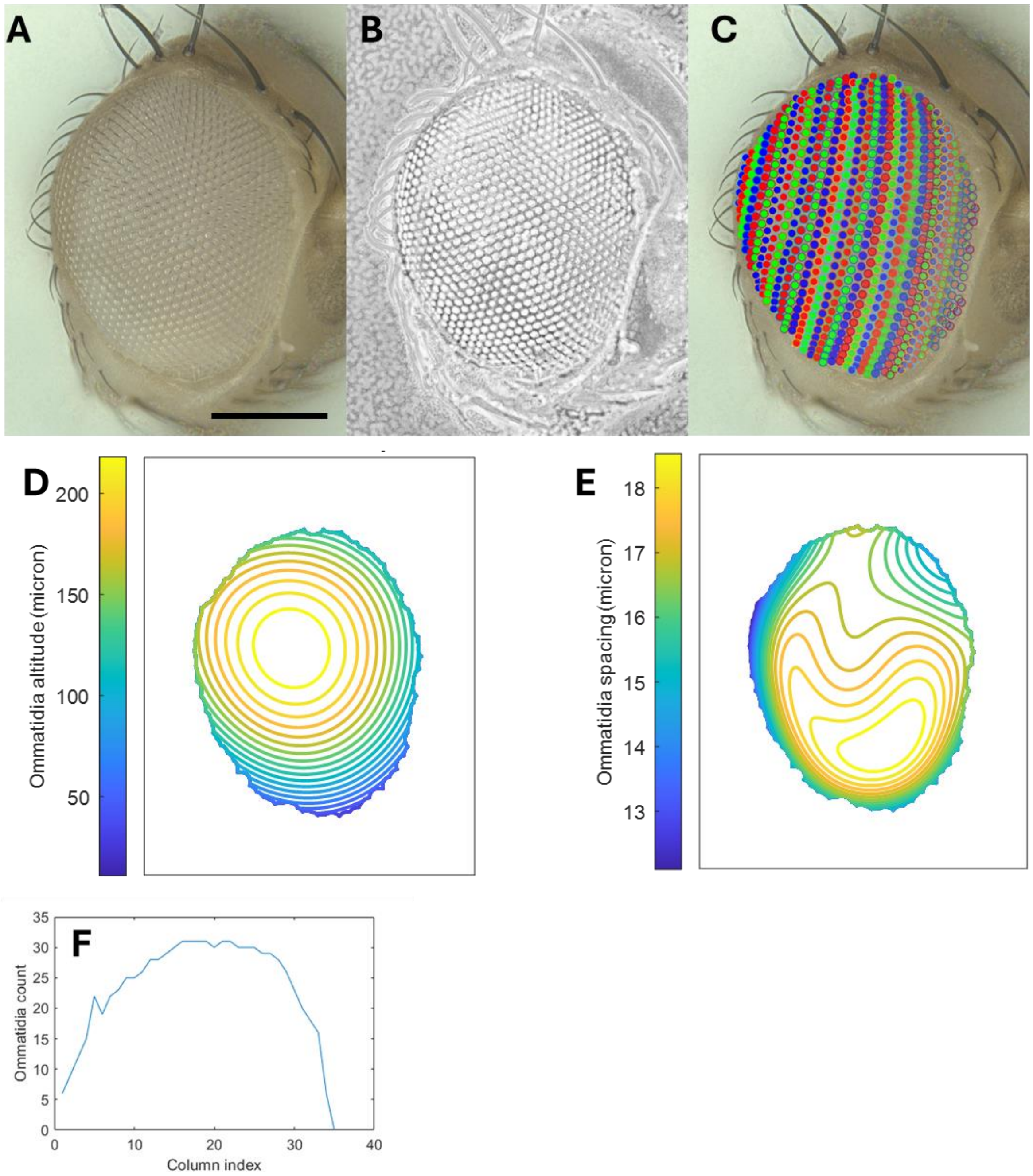


Figure S6. EyeHex morphological analysis on a brightfield image of a WTJ2 eye.

#### 3. Average ommatidia spacing profile

**Figure S7. Contour map of the average ommatidia spacing in WTJ2 eyes** (data from *Ramaekers et al., 2019*)

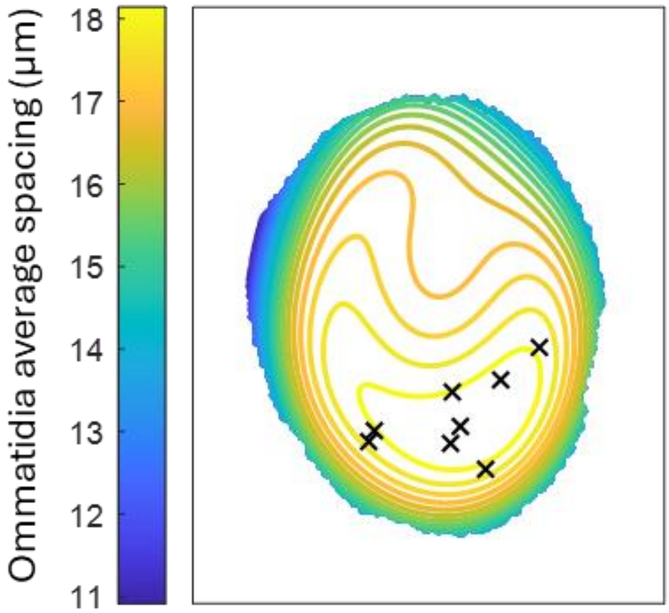


Figure S7. Contour map of the average spacing between adjacent ommatidia (in µm) projected onto the eye in Fig. S6. Black crosses indicate the locations of greatest inter-ommatidial distance (largest ommatidia) for individual WTJ2 eyes; data from n=8 eyes.
