## Supplementary file 2 for "EyeHex toolbox for complete segmentation of ommatidia in fruit fly eyes"

### Tran et al. Supplementary File 2

**Strain: Hikone-AS**

Scanning Electron Microscopy

Image from Ramaekers 2019

| | | EyeHex | | | | Manual counts | Elliptic estimation | | | |
|---|---|---|---|---|---|---|---|---|---|---|
| Index | **Image** | **pre-manual correction** | **post-manual correction** | **Correction%** | **Error%** | **Ramaekers 2019** | **DV rows** | **AP rows** | **ellipse area** | **side (Left/Right)** |
| 1 | 20100309-Hikone1s.tif | 776 | 713 | 8.84 | 0.00 | 713 | *29* | *32* | 728.85 | R |
| 2 | 20100309-Hikone2-lefteyes.tif | 689 | 728 | 5.36 | 0.97 | 721 | *28* | *34* | 747.70 | L |
| 3 | 20100309-Hikone3-righteyes | 909 | 760 | 19.61 | 0.00 | 760 | *30* | *33* | 777.54 | R |
| 4 | hik2-01-rights | 795 | 769 | 3.38 | 0.39 | 766 | *31* | *33* | 803.46 | R |
| 5 | hik2-03-rights | 868 | 727 | 19.39 | 0.27 | 729 | *29* | *33* | 751.63 | R |
| 6 | hik2-04-rights | 953 | 750 | 27.07 | 0.13 | 749 | *30* | *33* | 777.54 | R |
| | **mean** | **831.67** | **741.17** | **13.94** | **0.30** | **739.67** | *29.50* | *33.00* | **764.45** | |
| | standard deviation | 96.77 | 21.79 | 9.44 | 0.37 | 21.76 | *1.05* | *0.63* | 26.75 | |

**Strain: melanogaster CS**

Scanning Electron Microscopy

Image from Ramaekers 2019

| | | EyeHex | | | | Manual counts | Elliptic estimation | | | |
|---|---|---|---|---|---|---|---|---|---|---|
| **Index** | **Image** | **pre-manual correction** | **post-manual correction** | **Correction%** | **Error%** | **Ramaekers 2019** | **DV rows** | **AP rows** | **ellipse area** | **side (Left/Right)** |
| 1 | 20100309-CS1-righteyes | 947 | 874 | 8.35 | 0.23 | 876 | *32* | *35* | 879.65 | R |
| 2 | 20100309-CS2righteyes | 998 | 800 | 24.75 | 0.99 | 808 | *31* | *34* | 827.81 | R |
| 3 | 20100309-CS3righteyes | 849 | 869 | 2.30 | 0.12 | 868 | *32* | *36* | 904.78 | R |
| 4 | 20100309-CS4-lefteyes | 950 | 864 | 9.95 | 0.47 | 860 | *32* | *35* | 879.65 | L |
| 5 | 20100309-CS5-righteyes | 739 | 827 | 10.64 | 0.48 | 831 | *32* | *34* | 854.51 | R |
| 6 | CS2-01-rights | 1054 | 839 | 25.63 | 0.47 | 843 | *31* | *35* | 852.16 | R |
| 7 | CS2-02-rights | 878 | 879 | 0.11 | 0.11 | 880 | *32* | *36* | 904.78 | R |
| 8 | CS2-03-rights | 565 | 897 | 37.01 | 0.11 | 896 | *33* | *36* | 933.05 | R |
| 9 | CS2-04-lefts | 750 | 871 | 13.89 | 0.11 | 872 | *32* | *36* | 904.78 | L |
| | **mean** | **858.89** | **857.78** | **14.74** | **0.41** | **859.33** | *31.89* | *35.22* | **882.35** | |
| | standard deviation | 174.05 | 30.06 | 12.07 | 0.30 | 27.36 | *0.60* | *0.83* | 33.08 | |

**Strain: melanogaster WT25174 - DGRP collection**

Scanning electron microscopy

Images from Ramaekers 2019

| | | EyeHex | | | | Manual counts | Elliptic estimation | | | | |
|---|---|---|---|---|---|---|---|---|---|---|---|
| **Index** | **file** | **pre-manual correction** | **post-manual correction** | **Correction%** | **Error%** | **Ramaekers 2019** | **DV rows** | **AP rows** | **ellipse area** | **side (Left/Right)** | |
| 1 | 25174_1_lefts | 820 | 798 | 2.76 | 0.00 | 798 | *32* | *34* | 854.51 | L | |
| 2 | 25174_2_rights | 979 | 805 | 21.61 | 0.25 | 803 | *32* | *34* | 854.51 | R | |
| 3 | 25174_3_rights | 849 | 707 | 20.08 | 0.28 | 709 | *30* | *32* | 753.98 | R | |
| 4 | 25174_4_lefts | 797 | 715 | 11.47 | 0.00 | 715 | *30* | *32* | 753.98 | L | |
| 5 | 25174_5_lefts | 803 | 717 | 11.99 | 0.14 | 718 | *30* | *32* | 753.98 | L | |
| 6 | 25174_6_rights | 795 | 743 | 7.00 | 0.00 | 743 | *30* | *33* | 777.54 | R | |
| 7 | 25174_7_rights | 998 | 814 | 22.60 | 0.00 | 814 | *32* | *34* | 854.51 | R | bad automatic segmentation |
| 8 | 25174_8_lefts | 831 | 737 | 12.75 | 0.00 | 737 | *31* | *32* | 779.11 | L | |
| | **mean** | **859.00** | **754.50** | **13.78** | **0.08** | **754.63** | *30.88* | *32.88* | **797.77** | | |
| | standard deviation | 82.12 | 44.15 | 7.13 | 0.12 | 43.40 | *0.99* | *0.99* | 48.06 | | |

**Strain: D. pseudoobscura 121**

Scanning Electron Microscopy

Image from Ramaekers 2019

| | | EyeHex | | | | Manual counts | Elliptic estimation | | | |
|---|---|---|---|---|---|---|---|---|---|---|
| Index | **Image** | **pre-manual correction** | **post-manual correction** | **Correction%** | **Error%** | **Ramaekers 2019** | **DV rows** | **AP rows** | **ellipse area** | **side (Left/Right)** |
| 1 | pse121_2_01_lefts | 1136 | 1160 | 2.07 | 0.17 | 1158 | *36* | *42* | 1187.52 | L |
| 2 | pse121_2_02_rights | 1096 | 1194 | 8.21 | 0.08 | 1195 | *36* | *43* | 1215.80 | R |
| 3 | pse121_2_03_rights | 1078 | 1159 | 6.99 | 0.00 | 1159 | *36* | *43* | 1215.80 | R |
| 4 | pse121_2_05_lefts | 1167 | 1162 | 0.43 | 0.17 | 1160 | *36* | *43* | 1215.80 | L |
| 5 | pse121_2_06_rights | 1031 | 1206 | 14.51 | 0.25 | 1203 | *37* | *43* | 1249.57 | R |
| | **mean** | **1101.60** | **1176.20** | **6.44** | **0.14** | **1175.00** | ***36.20*** | ***42.80*** | **1216.90** | |
| | standard deviation | 52.52 | 22.16 | 5.56 | 0.10 | 22.10 | *0.45* | *0.45* | 21.99 | |

**Strain: Hikone-AS**

Light Imaging with Keyence macroscope

Images : This study

| | | | EyeHex | | | elliptic estimation | | |
|---|---|---|---|---|---|---|---|---|
| **Index** | **Image** | **pre-manual correction** | **post-manual correction** | **Correction%** | **DV rows** | **AP rows** | **ellipse area** | **side (Left/Right)** |
| 1 | 210608_hik10 | 803 | 765 | 4.97 | *29* | *35* | 797.18 | L |
| 2 | 210608_hik12 | 780 | 745 | 4.70 | *29* | *34* | 774.40 | R |
| 3 | 210608_hik13 | 793 | 760 | 4.34 | *30* | *34* | 801.11 | L |
| 4 | 210608_hik14 | 800 | 765 | 4.58 | *30* | *33* | 777.54 | L |
| 5 | 210608_hik15 | 815 | 768 | 6.12 | *29* | *34* | 774.40 | R |
| 6 | 210608_hik16 | 785 | 757 | 3.70 | *30* | *33* | 777.54 | R |
| 7 | 210608_hik17 | 801 | 751 | 6.66 | *29* | *34* | 774.40 | L |
| 8 | 210608_hik19 | 786 | 767 | 2.48 | *29* | *35* | 797.18 | R |
| 9 | 210608_hik20 | 758 | 732 | 3.55 | *28* | *34* | 747.70 | L |
| 10 | 210608_hik21 | 756 | 737 | 2.58 | *29* | *34* | 774.40 | L |
| 11 | 210608_hik22 | 774 | 721 | 7.35 | *28* | *34* | 747.70 | L |
| 12 | 210608_hik23 | 800 | 764 | 4.71 | *30* | *34* | 801.11 | L |
| | **mean** | **787.58** | **752.67** | **4.64** | *29.17* | *34.00* | **778.72** | |
| | std | 18.17 | 15.62 | 1.50 | *0.72* | *0.60* | 18.30 | |

**F1 progeny from ey3.5G x Df(4)J2**

Light Imaging with Keyence macroscope

Images from Ramaekers 2019

| Index | Images | EyeHex<br>pre-manual correction | <br>post-manual correction | Correction% | elliptic estimation<br>DV rows | <br>AP rows | <br>ellipse area | side (Left / Right) |
|---|---|---|---|---|---|---|---|---|
| 1 | 20190128_WTJ2_03.tif | 731 | 820 | 10.85 | *31* | *34* | 827.81 | R |
| 2 | 20190128_WTJ2_04.tif | 814 | 773 | 5.30 | *30* | *33* | 777.54 | L |
| 3 | 20190128_WTJ2_11.tif | 878 | 771 | 13.88 | *32* | *33* | 829.38 | L |
| 4 | 20190128_WTJ2_13.tif | 831 | 742 | 11.99 | *31* | *33* | 803.46 | R |
| 5 | 20190128_WTJ2_18.tif | 893 | 771 | 15.82 | *31* | *33* | 803.46 | R |
| 6 | 20190128_WTJ2_19.tif | 990 | 813 | 21.77 | *32* | *35* | 879.65 | R |
| 7 | 20190128_WTJ2_20.tif | 927 | 823 | 12.64 | *30* | *36* | 848.23 | L |
| 8 | 20190128_WTJ2_21.tif | 898 | 799 | 12.39 | *31* | *33* | 803.46 | L |
| | **mean** | **870.25** | **789.00** | **13.08** | *31.00* | *33.75* | **821.62** | |
| | std | 73.32 | 27.17 | 4.64 | *0.71* | *1.09* | 29.80 | |
